# Transcriptional Profiling and Genetic Analysis of a Cystic Fibrosis Airway-Relevant Model Shows Asymmetric Responses to Growth in a Polymicrobial Community

**DOI:** 10.1101/2023.05.24.542191

**Authors:** Christopher A. Kesthely, Rendi R. Rogers, Bassam El Hafi, Fabrice Jean-Pierre, George A. O’Toole

**Author notes:** To whom correspondence should be addressed Department of Microbiology and Immunology Geisel School of Medicine at Dartmouth, Rm. 202 Remsen Building, 66 North College Street, Hanover, NH, 03755.

## Abstract

Bacterial infections in the lungs of persons with cystic fibrosis are typically composed of multispecies biofilm-like communities, which modulate clinically relevant phenotypes that cannot be explained in the context of a single species culture. Most analyses to-date provide a picture of the transcriptional responses of individual pathogens, however, there is relatively little data describing the transcriptional landscape of clinically-relevant multispecies communities. Harnessing a previously described cystic fibrosis-relevant, polymicrobial community model consisting of *Pseudomonas aeruginosa, Staphylococcus aureus, Streptococcus sanguinis* and *Prevotella melaninogenica*, we performed an RNA-Seq analysis to elucidate the transcriptional profiles of the community grown in artificial sputum medium (ASM) as compared to growth in monoculture, without mucin, and in fresh medium supplemented with tobramycin. We provide evidence that, although the transcriptional profile of *P. aeruginosa* is community agnostic, the transcriptomes of *S. aureus* and *S. sanguinis* are community aware. Furthermore, *P. aeruginosa* and *P. melaninogenica* are transcriptionally sensitive to the presence of mucin in ASM, whereas *S. aureus* and *S. sanguinis* largely do not alter their transcriptional profiles in the presence of mucin when grown in a community. Only *P. aeruginosa* shows a robust response to tobramycin. Genetic studies of mutants altered in community-specific growth provide complementary data regarding how these microbes adapt to a community context.

**Importance:** Polymicrobial infections constitute the majority of infections in the cystic fibrosis (CF) airway, but their study has largely been neglected in a laboratory setting. Our lab previously reported a polymicrobial community that can explain clinical outcomes in the lungs of persons with CF. Here we obtain transcriptional profiles of the community versus monocultures to provide transcriptional information about how this model community responds to CF-related growth conditions and perturbations. Genetic studies provide complementary functional outputs to assess how the microbes adapt to life in a community.

## Introduction

Cystic fibrosis (CF), caused by a mutation in the CF transmembrane conductance regulator (*CFTR*) gene (1), is the most common autosomal recessive genetic disease in Caucasian populations (2). Mutations in *CFTR* have been demonstrated to impair the function of many organ systems, including the lungs. One of the hallmarks of disease for persons with CF (pwCF) is a loss of chloride channel function, resulting in increased mucous viscosity, reduction in mucociliary clearance and accumulation of mucus on the epithelial surface of the airways (3–5). The reduction of mucus clearance in the lungs of pwCF provides an ideal environment for the growth of bacterial and fungal pathogens; these chronic infections increase inflammation, resulting in tissue damage and increased mortality rates (6, 7).

Chronic infections originate early in the lives of pwCF (8). Once infected, many of the microbes that colonize the lungs of pwCF transition from an acute infection to a chronic biofilm mode of growth (9–13). The development of chronic infections in pwCF is associated with poor clinical outcomes, with 80-95% of pwCF succumbing to respiratory failure associated with chronic bacterial infections and the associated airway inflammation (14).

Examination of human infections with microscopy revealed the polymicrobial nature of many infections (15–17). Despite early knowledge that many infections are associated with more than one microbe, most efforts have focused on the study and treatment of individual bacterial species (15). Research focusing on polymicrobial communities has found that when multiple bacterial genera/species are present in a community, there is a disconnect between culture-based antimicrobial susceptibility testing and clinical responses (18–20).

Recent work from our lab has attempted to reconcile our lack of knowledge regarding polymicrobial biofilms in the context of CF infections (21). Our work has focused on a new clinically-informed model community comprised of a set of four CF-relevant microbes: *Pseudomonas aeruginosa, Staphylococcus aureus, Streptococcus sanguinis* and *Prevotella melaninogenica*, grown in artificial sputum medium (ASM) under anoxic conditions, mimicking the nutritional environment found in the mucus associated with the lungs of pwCF (21–26).

In this study, we developed transcriptional profiles of the four constituent members of a four-microbe community in a clinically-relevant growth condition, both in monoculture, in the four-microbe community, and in response to several perturbations. This data set will serve as a resource for the community. A key finding of these analyses is the differential impact on the transcriptome for these microbes grown in monoculture versus co-culture. We also performed genetic screens showing that a microbe with little change in transcription in the community (*P. aeruginosa*) also yield few mutants with small effect sizes in the community context. In contrast, for two microbes showing marked changes in the community transcriptome, we identified multiple mutants, some with large effect sizes. The transcriptional and genetic studies support the conclusion that microbes display asymmetric responses to growth in a polymicrobial community.

## Results and Discussion

### Transcriptional profiling studies

We performed a series of transcriptional profiling studies to better understand genes differentially expressed in monoculture versus our previously reported, clinically-relevant multispecies community (24). We also assessed the transcriptional profile of the community upon treatment with tobramycin in fresh medium (using fresh medium as a control), in medium lacking mucin, and for a community including a common *P. aeruginosa* variant frequently detected in the CF airway, that is, a Δ*lasR* mutant. We previously reported that LasR defective strains show an increased tolerance to tobramycin in the mixed community (24).

To this end, we obtained transcriptional profiles of each of the four constituent species, *P. aeruginosa* PA14, *S. aureus* Newman, *S. sanguinis* SK36 and *P. melaninogenica* ATCC 25845, across the different growth conditions outlined above (**Table 1, Supplemental Tables 1-4**). Co-culturing these four microbes in ASM supplemented with mucin provides a means to mimic the nutritional, environmental and community composition common in the CF airway (23–26), providing an ideal in vitro model to examine the transcriptional landscape of these four microbes in CF-relevant growth conditions.

**Table 1.**
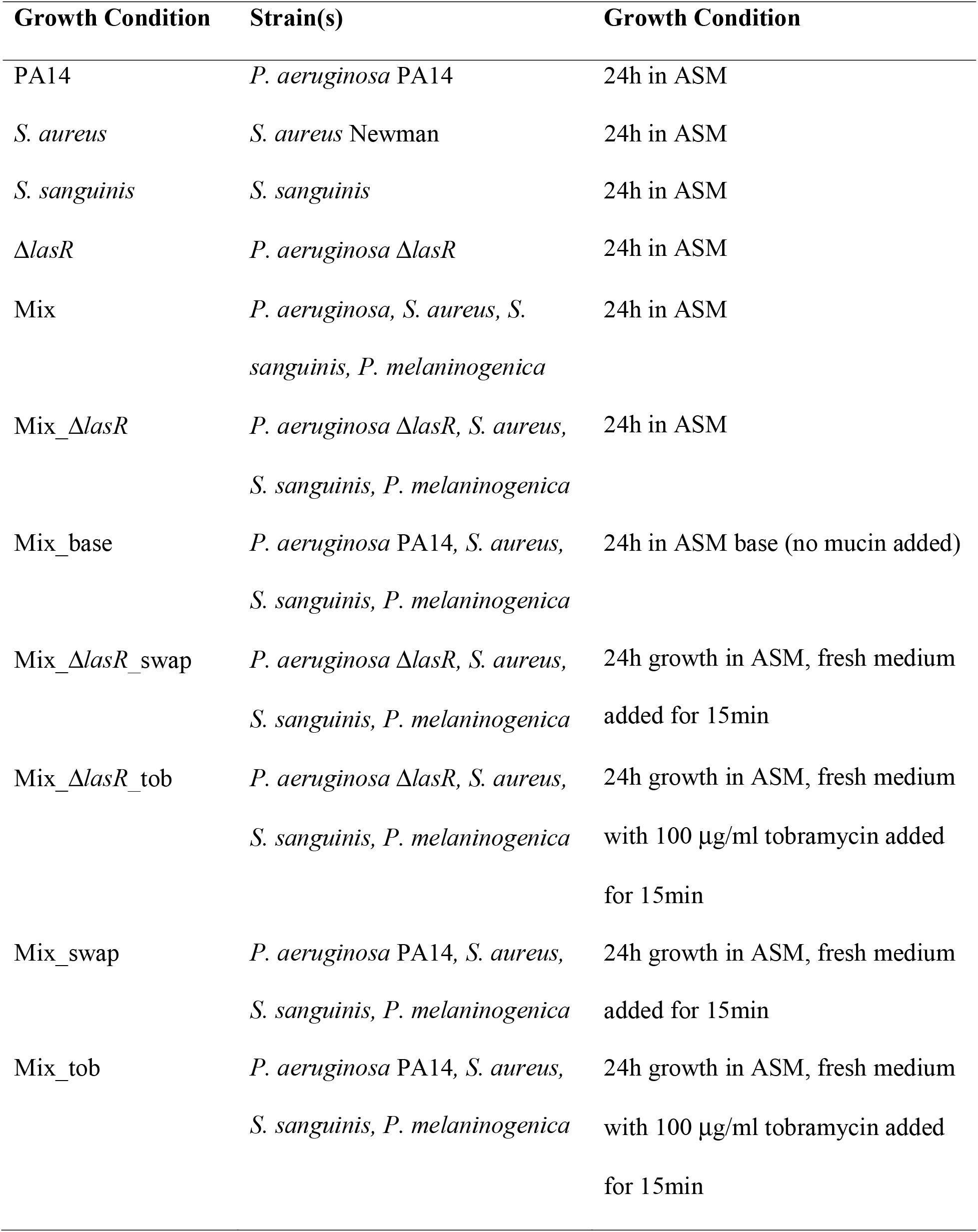
Labels for the strains and growth conditions used in the transcriptional studies.

### Response of the microbes to growth in the community

To begin analyzing our data, each genus’ RNA profiles in each of the indicated growth conditions were used to produce a series of PCA plots (**Figure 1 A-E**). Each point shown in the PCA plots represents the first two principal components for an individual biological replicate for the indicated condition, each with 4 replicates. We observed that *P. aeruginosa* and the Δ*lasR* mutant monocultures (**Figure 1A**, Mono and Δ*lasR*, respectively) clustered with their community-grown counterparts (**Figure 1A**, Mix and Mix_ Δ*lasR*, respectively). In contrast, the *S. aureus* monoculture (**Figure 1B**, Mono, pink) and *S. sanguinis* monoculture (**Figure 1C**, Mono, pink) were distinct from their corresponding community-associated counterparts (Mix, red). As previously reported, *Prevotella* does not grow in the model in monoculture (24) so there is no “*Prevotella* monoculture” condition (**Figure 1D**). The key for **Figures 1A-D** is shown in **Figure 1E**.

**Figure 1.**
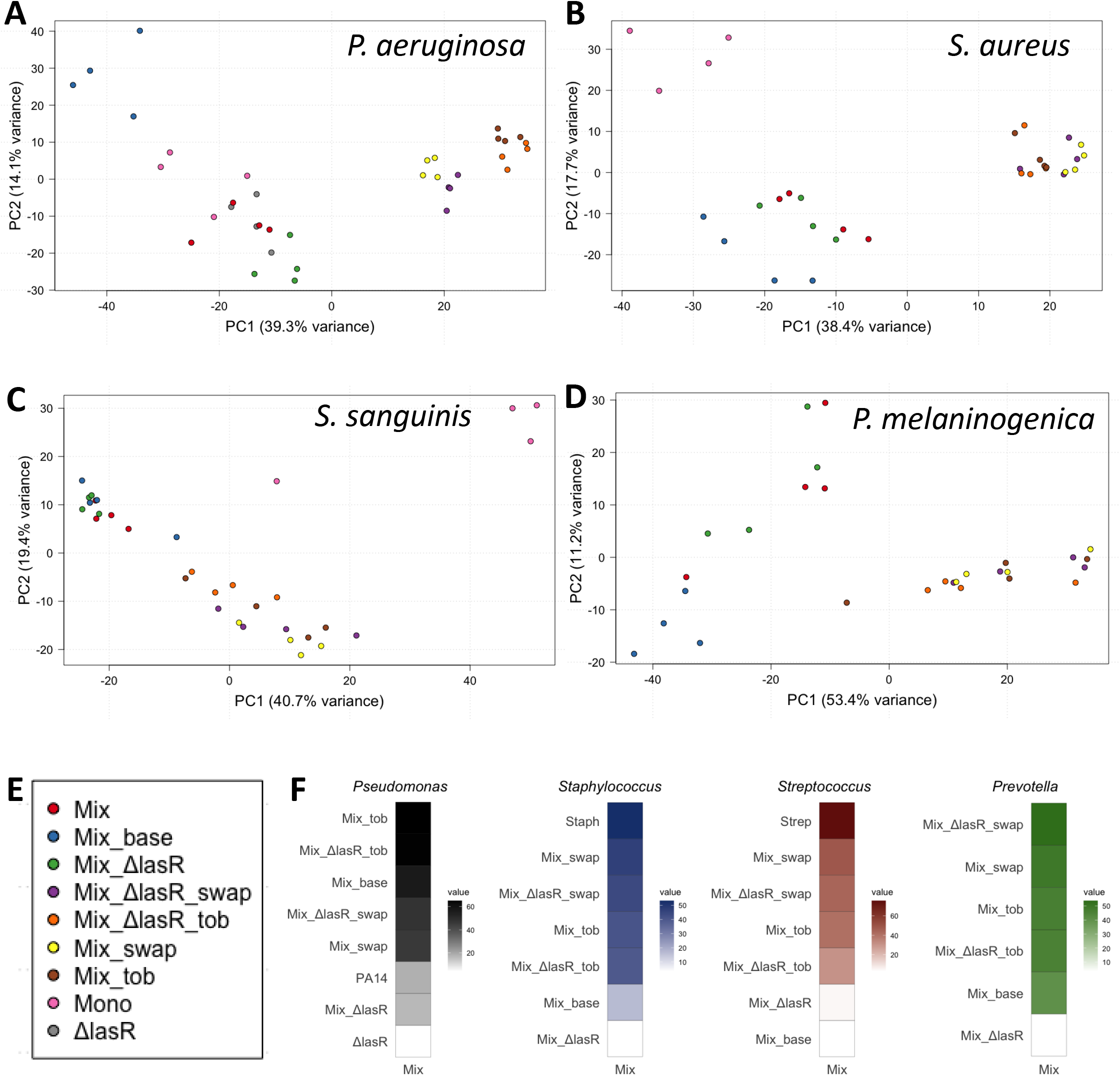
Variation in transcriptional profiles of monoculture versus the four-microbe community (Mix) in different culture conditions. A. PCA plot of *P. aeruginosa* transcriptional profiles across all culture conditions containing *P. aeruginosa.* B. PCA plot of *S. aureus* transcriptional profiles across all culture conditions containing *S. aureus.* C. PCA plot of *S. sanguinis* transcriptional profiles across all culture conditions containing *S. sanguinis.* D. PCA plot of *P. melaninogenica* transcriptional profiles across all culture conditions containing *P. melaninogenica.* The gene lists used to generate the diagrams in panels A-D can be found in Supplemental **Table 1**-4. E. Legend for the PCA plots in panels A-D. F. A distance matrix comparing the mixed community (Mixed) to the mean point for each of the other conditions (see **Table 1**) displayed in each PCA plot. Values were calculated via Euclidean distance between means and are included in the respective value marker present with each distance heatmap.

We performed a PERMANOVA on the data associated with each of the PCA plots to determine if the groupings were significantly different from each other. Pairwise tests demonstrated that only groupings within the *P. aeruginosa* PCA were classified as significantly different from each other. Specifically, while there was no significant difference for the WT or the *lasR* mutant in monoculture or the mixed community, or the Wt or *lasR* communities exposed to tobramycin, we did observe a significant difference when mucin was removed from the medium. For all other PCA plots shown in **Figure 1**, while we do see some significant changes in gene expression in response to perturbations, the overall transcriptional profiles are not statistically significant by PERMANOVA.

As a first step, the transcriptional data were compared using the mixed community (Mix) growth condition as a baseline to compare to the monoculture condition. *P. aeruginosa* wild type (WT) monocultures were found to have 19 downregulated genes and 135 upregulated genes when compared to the polymicrobial biofilm (**Supplemental Table 1**). Of the 135 genes upregulated in the *P. aeruginosa* WT monospecies biofilms, many of these genes are in pathways and/or operons implicated in biofilm formation and quorum sensing (**Supplemental Table 1)**.

*S. aureus* monocultures showed 50 downregulated genes and 159 upregulated genes when compared to *S. aureus* grown in the polymicrobial biofilm (**Supplemental Table 2**). Of the 50 downregulated genes, a large portion are implicated in metabolism and amino acid synthesis, while of the 159 upregulated genes, there is an enrichment for genes involved in *S. aureus* quorum sensing and thymidine metabolism (**Supplemental Table 2**).

*S. sanguinis* monocultures showed 240 downregulated genes and 211 upregulated genes when compared to *S. sanguinis* grown in the polymicrobial biofilm (**Supplemental Table 3**). The 240 downregulated genes were enriched for pathways involved in amino acid synthesis and D-xylulose-5-P synthesis, while the 211 upregulated genes were enriched for pathways involved in catabolism and metabolism (**Supplemental Table 3**).

Although the *S. aureus, S. sanguinis* and *P. aeruginosa* cultures appear to have a similar number of differentially regulated genes, the proportion of differentially regulated genes, in relation to genome size and number of genes, is less by a factor of two for *P. aeruginosa* compared to either *S. aureus* or *S. sanguinis*. Additionally, a closer examination of the magnitude of differential gene expression between the monocultures and co-cultures were lower for *P. aeruginosa* than for *S. aureus* or *S. sanguinis*. *P. aeruginosa* monocultures differed from the multispecies cultures by a log_2_FC of -2.87 – 5.32, the magnitude of differential gene expression of *S. aureus* monocultures differed from the multispecies cultures by a log_2_FC of -7.55 – 9.38 and *S. sanguinis* cultures differed by a log_2_FC of -11.10 – 7.56.

To further compare the transcriptional differences between the various conditions used in this study, we produced a distance matrix comparing the mixed community (Mix) to the mean point for each of the other conditions displayed in each PCA plot (**Figure 1F**). This distance matrix reinforces the finding that that *P. aeruginosa* (WT and the Δ*lasR* mutant) shows a modest shift in overall transcriptional profile when grown in monoculture versus mixed culture. Thus, it appears that *P. aeruginosa* is somewhat agnostic, in terms of gene expression, to monoculture growth conditions versus growth in the polymicrobial community. In contrast, the relative shift in transcriptional profiles for *S. aureus* and *S. sanguinis* in monoculture is much more striking when compared to growth in the mixed community (**Figure 1F**).

### Impact of fresh medium and tobramycin on transcriptional profiles

In a previous study, we found that the polymicrobial biofilm community conferred tolerance to tobramycin for biofilm-grown *S. aureus* and *S. sanguinis* compared to their planktonic counterparts (24). For *S. aureus,* growth in the mixed community conferred a modest (1-log) higher viability when exposed to tobramycin, while for *S. sanguinis*, growth in a mixed community resulted in 8-log greater viability than in the pure culture when both were exposed to tobramycin. In contrast, there was a statistical increase in the viability of the *S. sanguinis,* and no change to *S. aureus* when grown in mixed community when compared to the monoculture in the absence of tobramycin (24). Further, community-grown *P. melaninogenica* biofilms were unaltered by the presence or absence of tobramycin in the mixed community (**Supplemental Table 4**) (24). Unexpectedly, our previous work found a decrease in viability of biofilm-grown *P. aeruginosa* in the mixed community following the addition of tobramycin; this phenotype was reversed for the *P. aeruginosa lasR* mutant strain (24). We explore the impact of the *lasR* mutation below.

To better understand the basis of these changes in tolerance for microbes grown in the absence versus presence of the other community members (24), we assessed the transcriptional profiles of each of the constituent members of the four-microbe community upon the addition of fresh medium with tobramycin or fresh medium alone as the control. To assess the acute effects of addition of tobramycin, we harvested RNA 15 minutes after antibiotic (or medium control) exposure. We hypothesized that a 15-minute incubation would be sufficient to alter the transcriptional profiles without a significant loss in bacterial viability, which we confirmed experimentally (**Supplemental Figure 1**). The relatively short turnover time of mRNAs in bacteria (27, 28) supports our ability to investigate the acute effects of exposure to tobramycin.

We analyzed the response to acute tobramycin exposure versus the medium control in two ways. First, we generated a series of heatmaps to visualize the 50 most variably expressed genes (**Figure 2**) across all growth conditions in each tested species (**Table 1**). The list of these 50 genes is shown in **Supplemental Table 5**. The differentially expressed genes for the tobramycin exposure (Mix_tob, in brown) and the medium control (Mix_swap, in yellow, because the medium was “swapped”) formed a tight cluster for WT *P. aeruginosa* and for the *lasR* mutant, discrete from all other conditions– all other conditions were grown for 24 hours without the addition of fresh medium (**Figure 2A**). The tight association of the profiles of the top differentially expressed genes for the tobramycin exposure (Mix_tob) and the medium control (Mix_swap) are also reflected in the tight clustering of the respective samples in the *P. aeruginosa* PCA plot (**Figure 1A**).

**Figure 2.**
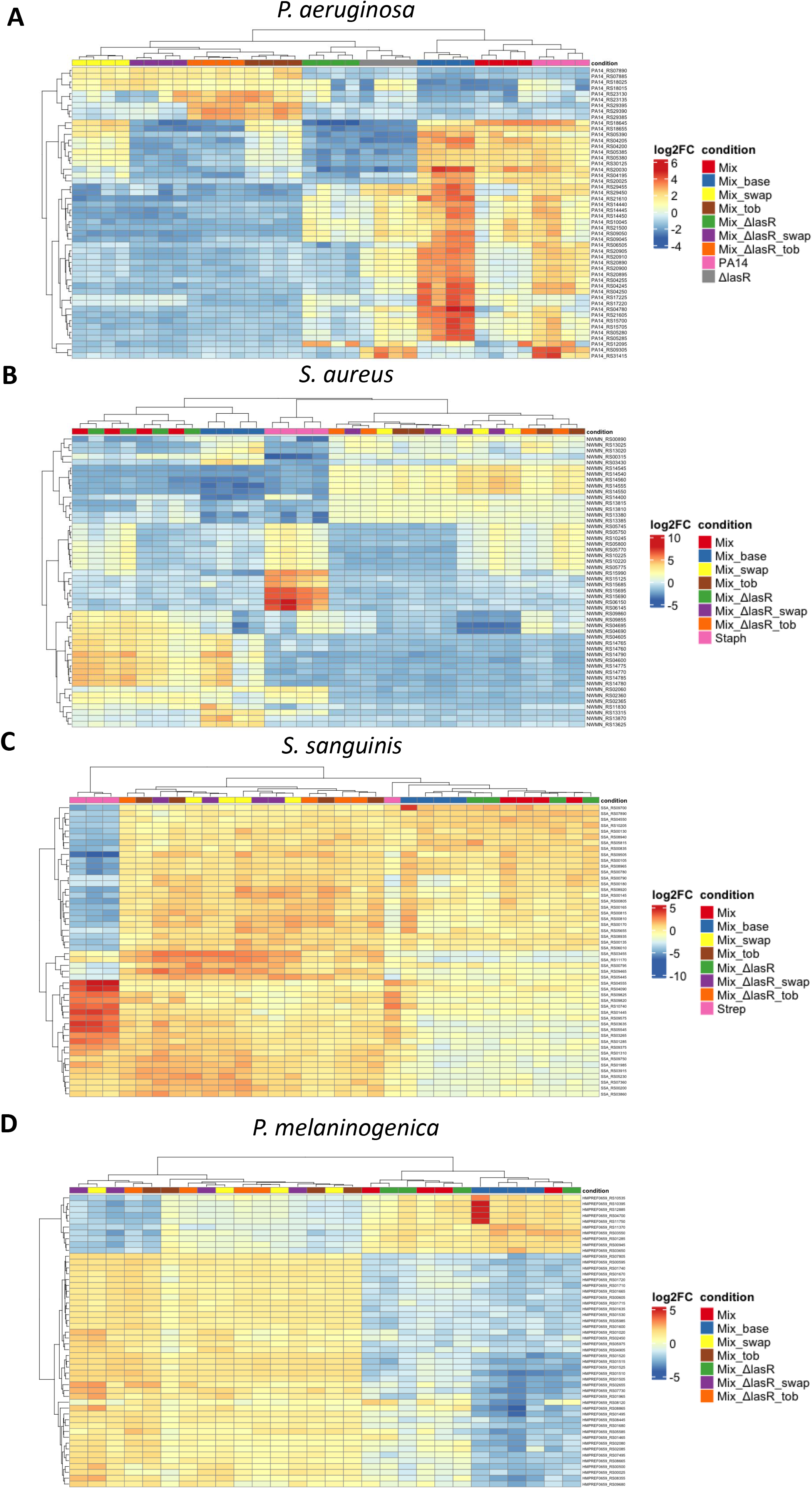
Heatmaps depicting the 50 most differentially expressed genes for each of the four bacteria present in monoculture and mixed communities in the different culture condition. **A**. The top 50 genes that are differentially expressed among the 9 different *P. aeruginosa* growth conditions. **B**. The top 50 differentially expressed genes in the 8 different *S. aureus* growth conditions. **C**. The top 50 differentially regulated genes in the 8 different *S. sanguinis* growth conditions. **D**. The top 50 differentially regulated genes in the 7 different *P. melaninogenica* growth conditions. Variation in gene expression was calculated with the rowVars function in the matrixStats package in R. The mean expression across all growth conditions was calculated, and the difference between the individual growth condition and the mean was utilized to produce a heatmap depicting the most differentially expressed genes across our various growth conditions through the use of the pheatmap package in R. The gene lists used to generate these diagrams can be found in **Supplemental Table 5**.

To further illustrate the influence of fresh medium +/- tobramycin on the mixed communities, a series of Venn diagrams were created to show the numbers of genes differentially expressed in these conditions compared to the mixed cultures at 24 hours (**Figure 3**). To generate the gene lists used to produce the Venn diagrams (**Supplemental Tables 6-9**), the RNA-Seq data were filtered to select only genes with a |log_2_FC| > 1.5 and an adjusted p-value < 0.05, then comparing samples exposed to only fresh medium to those exposed to fresh medium plus 100 μg/ml tobramycin.

**Figure 3.**
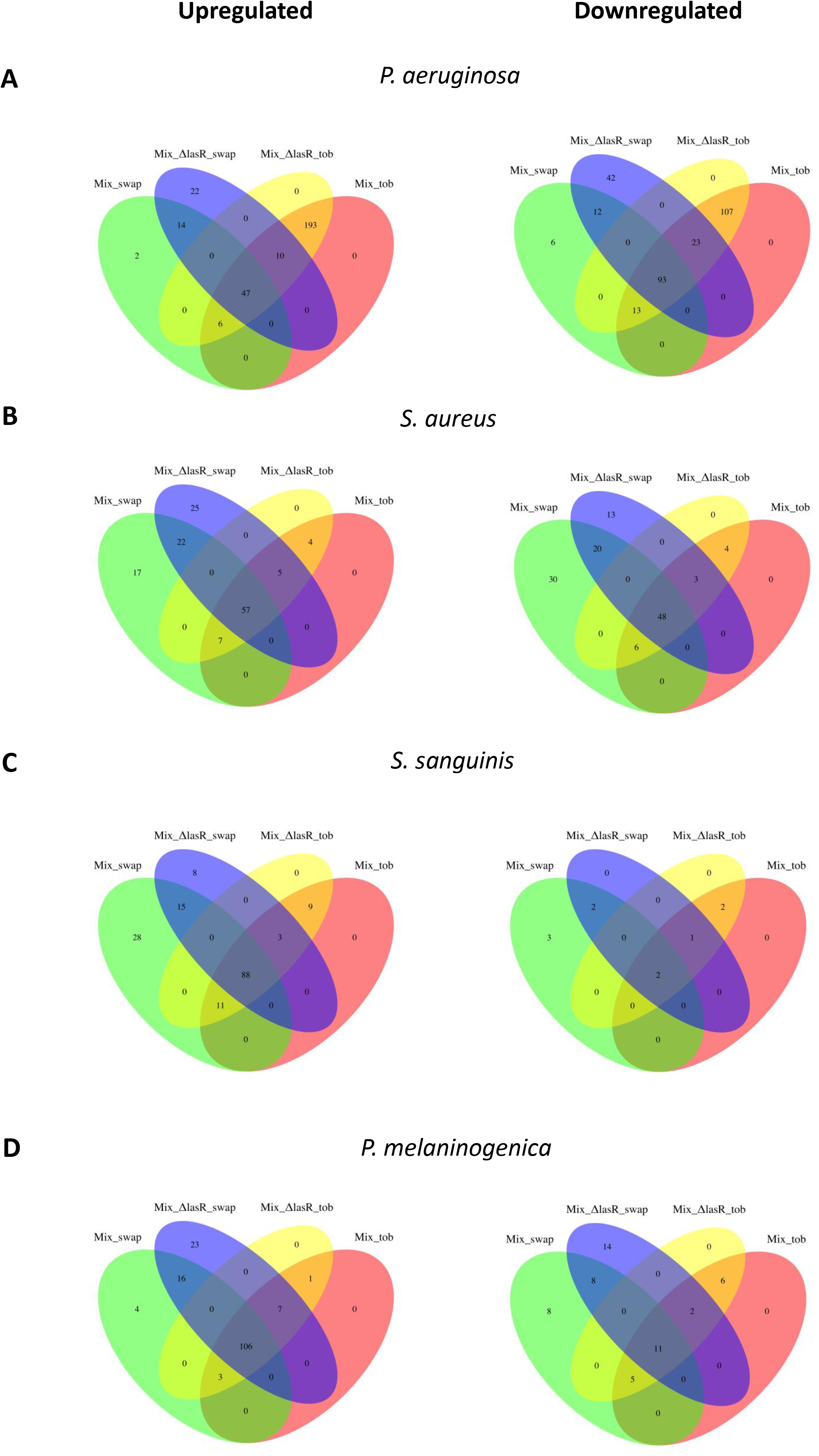
Genes differentially expressed from the mixed community (Mix) growth condition. Venn Diagrams were created to depict the number of genes differentially regulated in each of the growth conditions when compared to the mixed community grown in ASM (Mix) for **A**. *P. aeruginosa*, **B**. *S. aureus*, **C**. *S. sanguinis* and **D**. *P. melaninogenica*. To be included in the plot, genes had to be statistically differentially expressed (p < 0.05) and have a log_2_ fold change greater than or equal to the absolute value of 1.5. To provide a more informative figure, genes that were up and downregulated in comparison to the Mix biofilm condition were separated into separate Venn diagrams, left and right, respectively. Further analysis focused on the intersection of all datasets to elucidate the influence of fresh medium, and the intersection of solely the tobramycin conditions to elucidate the influence of tobramycin on the biofilm. The gene lists used to generate these diagrams can be found in **Supplemental Tables 6-9**.

Each of the Venn diagrams in **Figure 3** consist of 15 different regions. For *P. aeruginosa*, this analysis revealed a total of 47 upregulated and 93 downregulated genes associated with addition of fresh medium, and an additional 193 upregulated and 107 downregulated genes in response to tobramycin treatment (**Supplemental Table 6**). These differentially regulated genes were dominated by genes implicated in various metabolic pathways. Thus, *P. aeruginosa* showed a robust transcriptional response to the addition of tobramycin independent of the effect of adding fresh medium in the mixed community (**Figure 1F**).

The three other constituent members of the polymicrobial community showed a minimal response to the presence of tobramycin in the heatmap analysis of the top 50 genes, but their response to the addition of fresh medium was robust (**Figure 1B-F**, **Figure 2B-D, Supplemental Table 7-9**). This observation is best illustrated in **Figure 2B-D** by the co-clustering of samples exposed to fresh medium and fresh medium plus tobramycin for the WT and Δ*lasR* mutant communities. In contrast, for *P. aeruginosa* and as detailed above, the samples from cultures exposed to fresh medium formed a discrete cluster from those cultures exposed to fresh medium plus tobramycin (**Figure 2A**). The lack of a transcriptional response for *S. aureus* and *P. melaninogenica* is consistent with our previous report that these microbes do not show a robust change in the viability of biofilm-grown bacteria in monoculture versus a mixed community when exposed to tobramycin (24).

Interestingly, despite its lack of transcriptional response to tobramycin treatment, *S. sanguinis* biofilms treated with tobramycin do show a large and significant shift in viability when grown as a monoculture as compared to a mixed community (24). This change in behavior may be due to non-transcriptional mechanisms and/or response(s) to tobramycin that occur after the 15-minute exposure used in this study, or may be due to an unknown polymicrobial interaction protecting the *S. sanguinis* from tobramycin. The differential expression of genes in *P. aeruginosa,* but not the other three species in the polymicrobial community upon the addition of tobramycin to the cultures is consistent the modest efficacy of tobramycin in removal of *Pseudomonas* from the lungs of pwCF, however, does not provide insight into to reducing the remaining community members.

The transcriptional response to the exposure to fresh medium, independent of the exposure to tobramycin, for *S. aureus, S. sanguinis* and *P. melaninogenica*, is illustrated by the separation of the datapoints for these three microbes in the PCA plots in **Figure 1**, Venn diagrams in **Figure 3** and **Supplemental Tables 7-9**. As expected, the addition of fresh ASM, whether with or without tobramycin, drastically altered the transcriptional profiles of each species in the community (29–31). An analysis of the transcriptional responses of each of these species upon the addition of fresh medium failed to uncover any specific pathways. That is, many of the differentially regulated genes belonged to the “carbon metabolism” or “microbial metabolism in diverse environments” pathways as assessed by KEGG pathway analysis.

### Impact of the *lasR* mutation on transcriptional profiles

As mentioned above, we previously showed a substantial decrease in viability of biofilm-grown *P. aeruginosa* in the mixed community following the addition of tobramycin; this phenotype was reversed for the *P. aeruginosa lasR* mutant strain (24). Therefore, we included the Δ*lasR* mutant in several conditions: in monoculture, in the mixed community, in the mixed community exposed to fresh medium +/- tobramycin.

For *P. aeruginosa*, as expected, the top differentially expressed genes for the WT and the Δ*lasR* mutant formed discreet clusters (**Figure 2A**), whether grown in monoculture or in the mixed community for 24 hrs or exposed to fresh medium +/- tobramycin.

In contrast, *S. aureus* mixed cultures or the cultures exposed to fresh medium +/- tobramycin were unaffected by the *P. aeruginosa* Δ*lasR* mutant, which can be observed in PCA plot in **Figure 1B** and by the overlap of the most variably expressed genes in **Figure 2B**. Similarly, *S. sanguinis* appears to be agnostic to the presence of a *P. aeruginosa* Δ*lasR* mutant as the transcriptional profile of this microbe in the mixed community with WT *P. aeruginosa* and mixed community with the Δ*lasR* mutant are co-mingled in PCA plots (**Figure 1C**) and by clustering (**Figure 2C**). Finally, the Δ*lasR* mutant does not appear to impact the transcriptome of *P. melaninogenica,* as the profiles of this anaerobe grown with the WT and Δ*lasR* mutant are intermixed in the PCA plot (**Figure 1D**) and by clustering (**Figure 2D**). Overall, while loss of LasR function impacts the transcriptome of *P. aeruginosa*, as expected (32–34), mutating this quorum sensing system has a minimum impact on the other microbes in the community in the time frame used in our study.

### Impact of mucin on transcriptional profiles

The polymicrobial biofilms of both *P. aeruginosa* and *P. melaninogenica* are impacted by the presence or absence of mucin, while the *S. aureus* and *S. sanguinis* cultures show a minimal response to the presence/absence of mucin. Previous work demonstrated that bacteria like *P. melaninogenica* play a large role in mucin degradation (35, 36). Even though *S. sanguinis* has previously been shown to play a role in mucin degradation (35, 37, 38), we did not observe a transcriptional response indicating differential gene regulation when comparing Mix_base (no mucin) growth condition to the Mix growth condition (i.e., with mucin added).

As mentioned above, mucin degradation is associated with fermentative anaerobes such as *P. melaninogenica* (35). The fermentation of mucin by *P. melaninogenica* generates acetate and propionate, on which *P. aeruginosa* is partially dependent for growth in bacterial communities that mimic the lower airway of pwCF (35). In this work, the downregulation of *prpR,* which codes for the propionate catabolism regulator, in *P. aeruginosa* grown in the absence of mucin points to the bioavailability of propionate produced by *P. melaninogenica* in the presence of mucin. Likewise, a decrease in *P. melaninogenica* HMRPREF0695_A6215 transcription, coding for acetate kinase, which is responsible for the production of acetate and propionate, in the Mix_base (-mucin) community when compared to the Mix community (+mucin) points to a change in the metabolic profile of *P. melaninogenica* when grown in the mixed community in the absence of mucin.

### Summary of transcriptional responses

To further explore the transcriptional variability in each of the strains in the community, and across each growth condition, an upset plot was created for each bacterial species to compare the number of co-regulated genes across each of the growth conditions (**Table 1**) using the mixed community (Mix) as the comparison condition (**Figure 4**). The large number of co-regulated genes that were differentially regulated in similar conditions across each of our upset plots demonstrates the robustness of our dataset. To visualize the changes in how the transcriptional profiles changed across the various growth conditions, up and downregulated genes were separated to better understand how the transcriptional profiles of each organism was affected by the various growth conditions tested herein.

**Figure 4.**
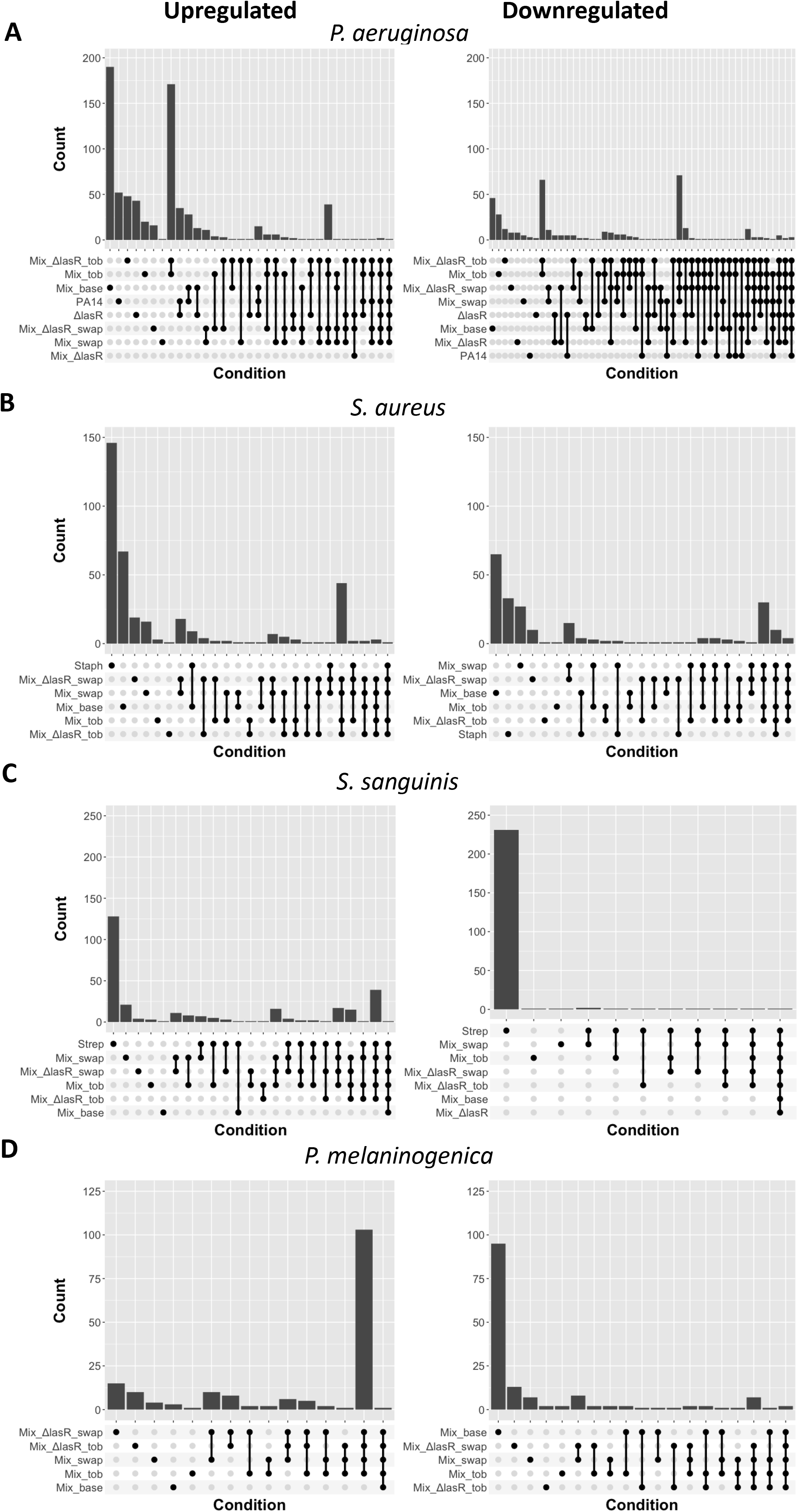
Upset plots to depict the number of differentially regulated genes. Upset plots were created to depict the number of genes differentially regulated in each of the growth conditions for both up and downregulated genes when compared to the mixed community grown in ASM (Mix). All genes used in the production of this plot were statistically differentially expressed (p < 0.05) with a log_2_ fold change greater than or equal to the absolute value of 1.5.

**Figure 4** demonstrates the roles that mucin and fresh medium have on the communities. With the exception of *S. sanguinis,* the lack of mucin produced the most differentially regulated genes in each of the other species across all of the treatments tested. The complexity of the mucin may play a role in secondary metabolism in each of these other microbial species. Since the human host produces 22 different mucins, it is also possible that the composition of the available mucin could play a role in the metabolism and interspecies interactions between each of the bacteria in our polymicrobial model (39, 40). Fresh medium, as described above, also plays a large role in the transcriptional profiles of each member of the tested polymicrobial communities. The large number of differentially regulated genes in the samples containing fresh medium support the ability of these bacteria to rapidly alter their transcription in responses to the presence of fresh nutrients.

### How genetic studies correlate with the transcriptional analyses

To complement the transcriptional analysis presented here, we performed genetic screens to ascertain the genes that, when mutated, would reduce survival in the community compared to monoculture, and at what magnitude (**Figure 5**). Genetic screens were performed with *P. aeruginosa, S. sanguinis and S. aureus* mutant libraries (41–43), and we compared viability of each mutant (via CFUs) when grown in polymicrobial community versus in monoculture.

**Figure 5.**
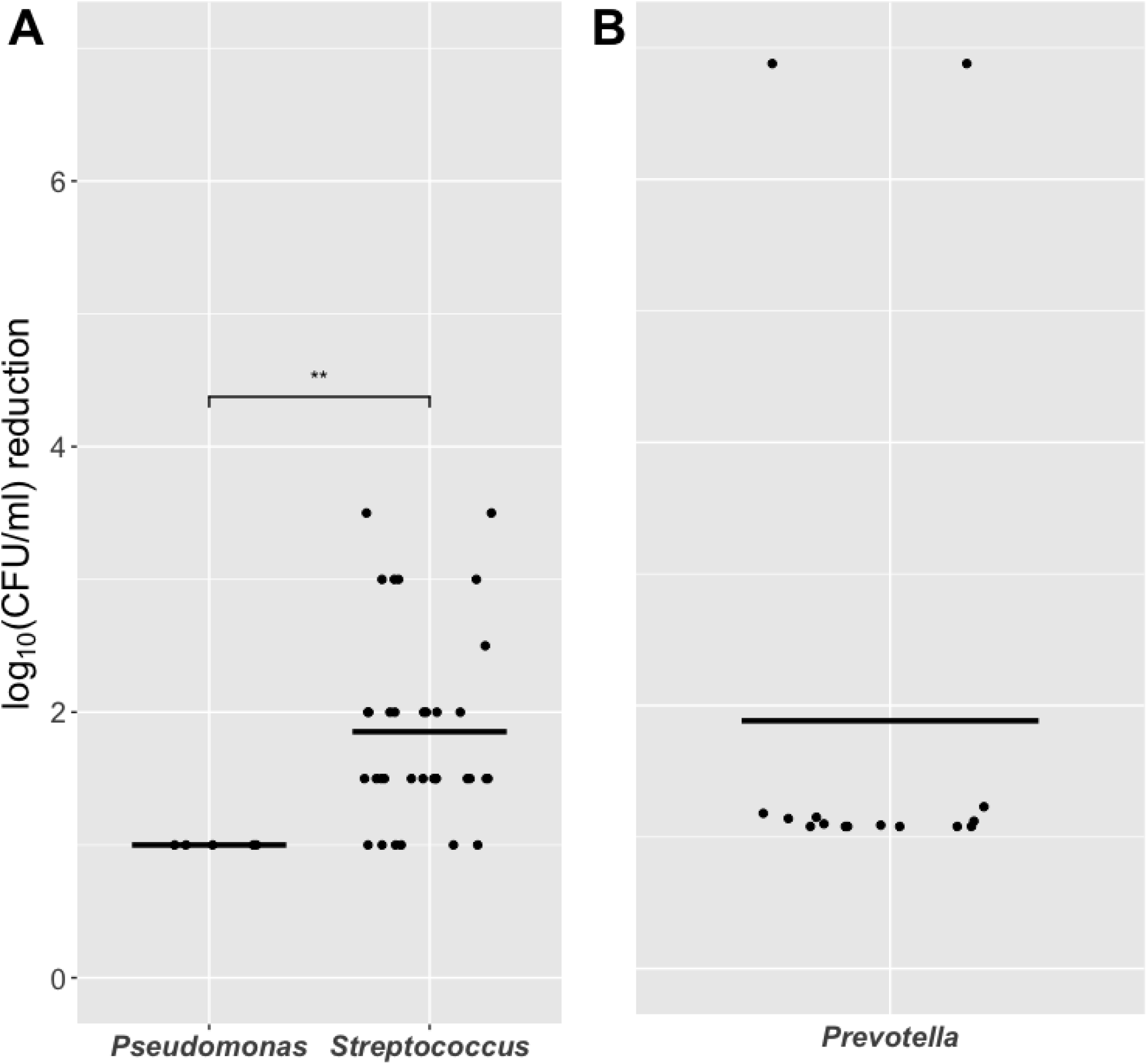
Fold change of mutants identified in genetic screens. Genetic screens using non- redundant transposon libraries were used to screen for mutations that reduced the growth of microbes in the polymicrobial community versus monoculture. **A**. Mutants that impacted the growth if *P. aeruginosa* (left) and *S. sanguinis* (right) when grown in the four-microbe community compared to monoculture. **B**. Mutants of *P. aeruginosa* that altered the growth of *P. melaninogenica* when grown as a co-culture. To be included, mutants had to decrease the CFUs of the strain from those grown in the monoculture, **A,** or when compared to *P. melaninogenica* grown in the presence of WT *P. aeruginosa* by ≥1-log. **B.** All mutants here showed no defect in monoculture growth (not shown). Stats were performed using both a wilcox test and with the ggsignif package in R. p=0.0078.

For the full *P. aeruginosa* transposon library (∼4500 mutants in non-essential genes), only 5 mutants were isolated; all of these strains resulted in a modest 1-log loss in CFU in the polymicrobial community as compared to when the mutant was grown in monoculture.

Following the same procedures, screening the whole *S. sanguinis* transposon library (∼2100 mutants) yielded 34 genes with reduced CFU when mutants were grown in a mixed community compared to a monoculture. When compared with the five genes that restricted the growth of *P. aeruginosa* in the community, these 34 genes produced a significantly larger log-fold change (**Figure 5A**, right side), varying from 1-log to 3.5-log fold change in viability.

Interestingly, screening half of the *S. aureus* Newman transposon library (∼1500 mutants) did not yield mutants with reduced viability when grown in the community.

Due to the lack of genetic tools in *P. melaninogenica,* and its inability to grow as a monoculture in ASM, we focused on *P. aeruginosa* mutants that limited the growth of *P. melaninogenica* in co-culture compared to co-culture with WT *P. aeruginosa*. The genetic screen of ∼4500 mutants from the *P. aeruginosa* mutant library identified 15 *P. aeruginosa* mutants that impacted the growth of *P. melaninogenica* (**Figure 5B)**. Thirteen of the *P. aeruginosa* mutations provided ∼1-log fold change in the growth of *P. melaninogenica,* while two mutants abolished the ability of *P. melaninogenica* to grow in co-culture conditions. All of the transposon mutants identified here grew as well as the WT strain in mono- and co-culture.

Together, the genetic analysis largely mirrors the transcriptional study, that is, *P. aeruginosa*, which is agnostic to community growth yielded few mutants with small effects on viability in the community. In contrast, for *S. sanguinis* and *P. melaninogenica*, which showed a more robust transcriptional response to growth in the community, also yielded more mutants with a larger range of phenotypes. The one exception was *S. aureus*, which despite its robust transcriptional responses across the various conditions analyzed here, failed to yield any mutants that showed a defect specific to the community, perhaps suggesting that *S. aureus* is well suited to compete against other microbes in a community via redundant systems not accessible to genetic screens.

## Conclusions

Our findings indicate that in the model polymicrobial community, the transcriptional profile of *P. aeruginosa* remains relatively unchanged by the presence of the other members of our community, a finding reflected by the genetic analysis. That is, both experimental approaches used here reiterate capacity of *P. aeruginosa* to thrive in the presence of other microbes across a range of environments. In contrast, the transcriptional profiles of the other members of the community are influenced by each other and environmental perturbations. Again, the transcriptional analysis is largely reflected by our genetic studies, with the exception of *S. aureus*.

While a great deal of work has focused on *P. aeruginosa* monospecies biofilms, and the role of the transcriptional regulator LasR impacting the transcription of hundreds of genes in this microbe (44–49), our findings indicate that a Δ*lasR* mutant of *P. aeruginosa* had a negligible influence on the transcriptional profiles of the other members of the community.

As expected, the addition of fresh medium, both with and without tobramycin, drastically altered the transcriptional responses of each of the members of the polymicrobial community. However, surprisingly, the addition of tobramycin only altered the acute transcriptional response of *P. aeruginosa.* The lack of a transcriptional response to tobramycin by the other members of this community confirms previous work performed in our lab demonstrating that upon tobramycin treatment, the viability of *S. sanguinis, S. aureus* and *P. melaninogenica* largely remain unchanged when grown in the polymicrobial community (24).

Here we have generated and performed an initial transcriptional analysis of a model polymicrobial community relevant to CF. We hope that providing the complete dataset to the broader research community will stimulate further hypotheses and help to advance the treatment of polymicrobial infections in CF-relevant infections. Due to the complexity of many polymicrobial infections, we believe that we have established a framework to further investigate polymicrobial infections using an approach that enables hypothesis generation, while simultaneously implicating specific genes and pathways for their roles in complex microbe-microbe interactions.

## Materials and Methods

### Bacterial strains and culture conditions

*Pseudomonas aeruginosa* PA14, *Staphylococcus aureus* Newman, *Streptococcus sanguinis* SK36 and *Prevotella melaninogenica* ATCC 25845 used in this study are listed in **Table 1**. All strains were cultivated as previously described (24). Briefly, *P. aeruginosa* and *S. aureus* cultures were grown in Tryptic Soy Broth (TSB) with shaking at 37°C. *P. melaninogenica* cultures were grown in TSB supplemented with 0.5% Yeast Extract (YE), 5 µg/mL hemin, 2.85 mM L-cysteine hydrochloride and 1 µg/mL menadione (*Prevotella* growth medium – PGM). *S. sanguinis* overnight cultures were grown in Todd-Hewitt Broth supplemented with 0.5% YE (THY) at 37°C with 5% CO_2_. Artificial sputum medium with (ASM) or without mucin (ASM base) was prepared as previously reported (24) and supplemented with 100 mM 3-morpholinopropane-1-sulfonic acid (MOPS) to maintain a pH of 6.80.

### Microbial growth assays

Culture experiments were performed as previously described by (24) with modifications. Briefly, cells from overnight liquid cultures of *P. aeruginosa*, *S. aureus*, *S. sanguinis* and *P. melaninogenica* were individually collected and washed twice (for *P. aeruginosa* and *S. aureus*) or once (for *S. sanguinis* and *P. melaninogenica*) in sterile Phosphate-Buffered Saline (PBS) by centrifuging at 10,000 x *g* for 2 minutes. After the final wash, cells were resuspended in ASM base. The optical density (OD_600_) was then measured for each bacterial suspension and diluted to an OD_600_ of 0.2 in either ASM or ASM base, depending on the subsequent growth condition. Monocultures and co-cultures were prepared from the OD_600_ = 0.2 suspensions and further diluted to a final OD_600_ of 0.01 for each microbe in ASM or ASM base, depending on the subsequent growth condition. Per each monoculture and co-culture conditions, a full polystyrene flat-bottom 96-well plate was inoculated with 100 µl of bacterial suspension. Plates were incubated using an AnaeroPak-Anaerobic container with a GasPak sachet (ThermoFisher) at 37 °C for 24 hours. After incubation, plates were processed and split into three separate groups. For group 1, which included monocultures and co-cultures communities grown in ASM or ASM base, unattached cells were aspirated with a multichannel pipette. Pre-formed biofilms were then washed once with sterile PBS and resuspended in 50 µl of RNAprotect (Qiagen) using a sterile 96-pin replicator. For the remaining groups (co-culture communities with WT or Δ*lasR P. aeruginosa*), plates were anaerobically transferred from an AnaeroPak-Anaerobic container with a GasPak sachet to an anoxic environmental chamber (Whitley A55 - Don Whitley Scientific, Victoria Works, UK) with 10% CO_2_, 10% H_2_, 80% N_2_ mixed gas at 37 °C. Unattached cells of co-culture communities grown in ASM were aspirated with a multichannel pipette and 100 µl of either fresh ASM (group 2) or ASM supplemented with 100 µg/mL tobramycin (group 3), was added to the pre-formed biofilms for 15 minutes.

After incubation, the supernatants were removed and the biofilms were washed once with sterile PBS. For all groups, a volume of 50 µl of RNAprotect was added to the biofilms and collected using a sterile 96-pin replicator. For all conditions, a control plate of the monocultures and co-cultures was 10-fold serially diluted and plated on selective media, as previously reported (24), as a control to confirm the expected viable counts of bacteria grown under each condition.

### RNA purification, sequencing and analysis

Following growth and collection of biofilms in RNAprotect, bacterial lysis and RNA extraction were performed following the protocols outlined in the RNeasy Mini Kit (Qiagen). Because of *S. aureus* in the mixed communities, cells lysis conditions included a combination of lysozyme (800 μg/ml) and lysostaphin (12.5 μg/ml). The concentration and quality of the RNA was checked on a Nanodrop 2000 (Thermo) prior to sending the samples for library preparation and sequencing (SEQCENTER). Briefly, samples were DNase treated (Invitrogen) prior to library preparation using the Illumina Stranded Total RNA Prep Ligation with Ribo-Qero Plus kit and 10 bp IDT for Illumina indices. Sequencing was performed on a NextSeq 2000 (2 x 51 bp reads). SEQCENTER performed demultiplexing, quality control and adapter trimming with bcl-convert (v3.9.3) (50). Upon obtaining the data, samples were subsequently analyzed with Cutadapt (51) to a phred score of 20, a minim length of 10 bp and using Cutadapt’s nextseq-trim function set at 10. Sequencing reads were mapped to the *P. aeruginosa* PA14 genome (NC_008463.1), *S. aureus* Newman genome (NC_009641.1), *S. sanguinis* SK36 genome (NC_009009.1) or the *P. melaninogenica* ATCC 25845 genome (NC_014370.1 and NC_014371.1) using bowtie2 (52) in paired end read using default parameters. The sam files were converted to sorted.sam files using samtools (53) prior to using featureCounts (54)with default parameters to calculate reads per gene. The differential expression of genes in each condition was determined through the use of DEseq2 (55). Read counts were normalized using the estimateSizeFactors function. Genes with read counts under 10 were dropped from our dataset. The biological coefficient of variance was estimated, and a linear model was fit to determine the differential expression of genes between samples. Our false discovery rate was adjusted from p-values using the Bonferroni correction (56). To compare the transcriptional profiles between samples, our datasets were further filtered to ensure that only genes with a differential expression by a log_2_ fold change (logFC) threshold greater than the absolute value of 1.5 and an adjusted p-value less than 0.05 were used in our downstream analyses. PCA plots were produced in base R, while heatmaps utilized the ComplexHeatmap (57) and pheatmap (58) R packages. Venn diagrams and upset plots were produced with the VennDiagram (59) and ggupset (60) R packages respectively.

### Genetic screens

For the genetic screens, we utilized non-redundant transposon mutant libraries for each of *P. aeruginosa* PA14, *S. sanguins* SK36 and *S. aureus* Newman (41–43). Overnight liquid cultures of mutant libraries were grown in polystyrene flat-bottom 96-well plates with each well containing a different mutant growing in 100uL of TSB for *P. aeruginosa,* THY for *S. sanguinis* and TSB+YE for *S. aureus* Newman. The 96 well plates containing monoculture conditions and co-culture conditions in ASM with *S. aureus, S. sanguinis,* and *P. melananogenica* were prepared using methods described above and inoculated with 1-2 uL of the *P. aeruginosa* PA14 (41), *S. sanguinis* (42) and *S. aureus* Newman (43) mutant libraries using a 96-pin replicator.

For all screens, plates were incubated using an AnaeroPak-Anaerobic container with a GasPak sachet (ThermoFisher) at 37 °C for 24 hours. After incubation, unattached cells were aspirated with a multichannel pipette. Pre-formed biofilms were then washed once with sterile PBS. Cells were then resuspended in 50 µl PBS and spotted onto the appropriate selective medium for each microbe using a sterile 96-pin replicator, as previously reported (24). Mutants that showed confluent growth in the monoculture condition but little to no growth in the mixed condition were selected as candidates for re-testing. For confirmation, each candidate was grown in triplicate in the monoculture and mixed conditions as described above. After incubation, all cultures were 10-fold serially diluted and plated on PIA for *P. aeruginosa,* SSA from *S. sanguinis,* TSB for *S. aureus, or* TSB+YE for *P. melaninogenica* to quantify cell counts by colony forming units (CFUs), as reported.

### Statistical Analyses

All statistical analyses performed in this paper were done in R. Principal component analyses (PCA) were performed on normalized and log transformed count data obtained from a DESeq2 analysis performed on the read counts for each of the 4 RNA-Seq replicates analyzed in this paper.

The adonis2 function from the package Vegan (61) was utilized to compare the overall dataset, and found a significant difference between each strain (p<0.001 for all four species tested). To examine pairwise groupings between the various strain groupings in the PCA plots (**Figure 1A-D**), we utilized the pairwise.adonis function from the package Vegan (61, 62).

Analyses were performed using the PC1 and PC2 data for each respective species, and comparisons made between each of the respective growth conditions. Both the adonis 2 and pairwise.adonis analyses utilized Euclidean distances as the “method”, or “sim.method” variable, respectively, to remain consistent with the Euclidean distances that were calculated to compare groupings in **Figure 1F**.

To compare the *P. aeruginosa* and *S. sanguinis* screen data, p-values were calculated through the use of the wilcox.test function, and the ggsignif package in R (63). Due to the difference in how the screen data were generated for *P. aeruginosa, S. aureus, S. sanguinis* and the *P. melaninogenica* samples, statistical analysis was not done to compare the *P. melaninogenica* data to the *P. aeruginosa* and *S. sanguinis* data.

## Data availability statement

Raw sequencing data was submitted as an SRA to NCBI and is available under the BioProject accession PRJNA971914. UNIX and R code used to produce the data and figures presented in this paper are available on GitHub (https://github.com/GeiselBiofilm).

## Acknowledgements

This work was supported by Cystic Fibrosis Foundation (JEAN21F0) to FJP and National Institutes of Health (R01 AI155424) to GAO. We would like to thank Dr. Tom Hampton and the DartCF core (NIH grant P30 KD117469 and CFF grant STANTO19R0) for conversations regarding data analysis.

