## Supplemental Figure 1 for "Transcriptional Profiling and Genetic Analysis of a Cystic Fibrosis Airway-Relevant Model Shows Asymmetric Responses to Growth in a Polymicrobial Community"

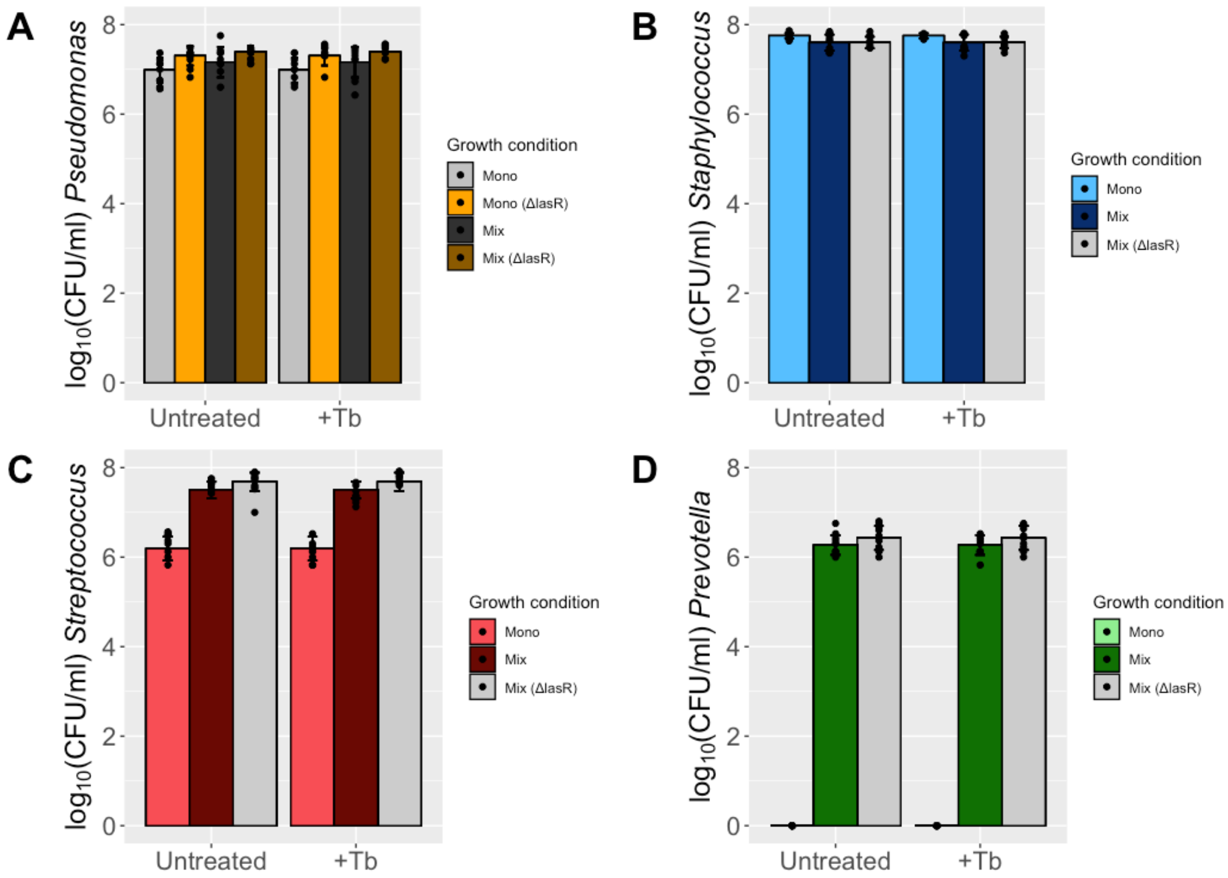


**Figure S1. Acute exposure to 100ug/ml tobramycin does not impact bacterial viability.** To confirm that viability was not impacted by an acute 15-minute exposure of tobramycin, CFU/ml counts of both monoculture and polymicrobial cultures exposed to fresh medium or fresh medium supplemented with 100μg/ml tobramycin were performed. **A**. Impact of tobramycin on *P. aeruginosa* mono and co-cultures. **B**. Impact of tobramycin on *S. aureus* mono and co-cultures. **C**. Impact of tobramycin on *S. sanguinis* mono and co-cultures. **D.** Impact of tobramycin on *P. melaninogenica* mono and co-cultures. CFU counts were not altered by the presence of 100 μg/ml tobramycin in any of the growth conditions. Three biological replicates with 3 technical replicates were performed, mean values were used to produce the bars, with standard deviation plotted as error. Individual datapoints are overlayed on the graphs. An ANOVA with a post hoc Bonferroni correction was performed on each of the datasets. For all datasets $p>0.05$.
